# Plaque-associated Microglial Polarization in Visual Brain Regions of the 5xFAD Mouse Model

**DOI:** 10.64898/2026.02.27.708522

**Authors:** Shaylah McCool, Jennie C. Smith, Matthew J. Van Hook

## Abstract

Alzheimer’s disease (AD), a neurodegenerative disorder associated with amyloid beta (Aβ) plaque deposition, leads to cognitive decline in affected individuals. Vision changes are some of the first reported symptoms in AD with studies showing both decline in functions performed by the visual system as well as associations between vision loss and cognitive impairment in AD patients. Due to the increasing number of individuals diagnosed with AD and its early impact on vision, we sought to provide an in-depth analysis using immunohistochemistry and 2-photon imaging techniques in the 5xFAD mouse model of amyloidosis to examine specifically how Aβ, a primary pathology typically preceding many other AD-associated pathologies, affects visual regions of the brain and how microglia, key immune regulators of the brain’s environment, respond to this AD-like pathology. We found that in the pathway for image-forming vision, including the dorsolateral geniculate nucleus (dLGN) and the primary visual cortex (V1), there was significant Aβ pathology and shifts in microglial morphology to an amoeboid state and increased phagocytic activity. However, in non-image-forming visual brain regions such as the superior colliculus (SC) and suprachiasmatic nucleus (SCN), there was minimal Aβ pathology, ramified microglial morphology, and minimal phagocytic activity. Overall, visual brain regions associated with Aβ plaque deposition experience significant microglial polarization when examining both morphology and function.

## 1. INTRODUCTION

Alzheimer’s disease (AD) is the most common form of dementia affecting the aging population with major pathologies including amyloid beta (Aβ) plaque and neurofibrillary tangle (NFT) formation followed by atrophy of the brain leading to memory and cognitive decline. Individuals with AD often report changes in vision, and the visual system has been studied as a potential early biomarker for AD (Javaid et al., 2016; Paik et al., 2020; Poudel et al., 2025). Both pathological changes – Aβ plaques and NFTs in visual brain regions, fewer optic nerve axons, reduced retinal thickness and vasculature – and visual manifestations of that pathology – oculomotor dysfunction, circadian rhythms disruptions, impaired multisensory integration, and decreased contrast sensitivity, color vision, visual acuity, and visual field – are documented in patients diagnosed with AD (Katz and Rimmer, 1989; Kusne et al., 2016). However, the mechanisms underlying immune responses to AD-like pathology in visual brain regions are poorly understood.

Microglia, the macrophages of the brain, are responsible for maintaining a healthy environment for the neurons, shifting their functional properties and morphology in response to neural trauma, pathological processes, and neurodegeneration (Wang et al., 2021). Using the 5xFAD mouse model, which develops Aβ plaques without the presence of NFTs, we aimed to examine the response of microglia to Aβ pathology in several visual brain regions including the dorsolateral geniculate nucleus (dLGN), the superior colliculus (SC), the suprachiasmatic nucleus (SCN), and the primary visual cortex (V1) (Oakley et al., 2006). Although these regions serve different purposes in the context of vision, the dLGN, SC, and SCN all receive direct synaptic input from retinal ganglion cells. The dLGN is a thalamic relay that connects the retina to V1 in the pathway for conscious vision, and although V1 does not receive direct input from the retina, it is reciprocally connected with the dLGN (Seabrook et al., 2017; Guido et al., 2018). The SC is a widely connected midbrain structure responsible for the integration of visual, auditory, and tactile stimuli and important for sensorimotor behaviors with its primary role in visual function being the direction of head and eye movements (Seabrook et al., 2017; Cang et al., 2018; Benevidez et al., 2021). The SCN, a small hypothalamic region, is responsible for setting circadian rhythms and receives input from intrinsically photosensitive retinal ganglion cells (Nassan et al., 2021).

In this study, we analyzed visual brain regions for AD-like pathology and microglial behavior because, in AD patients, Aβ plaques have been found in all four of these visual brain regions to varying degrees as well as disruption of systems regulated by these four regions including loss of different measures of visual function, altered circadian rhythms, and oculomotor dysfunction. As such, we hypothesized that Aβ plaques would have differential effects on microglial morphology and activity in visual brain regions responsible for diverse functions. Using immunohistochemistry and 2-photon imaging techniques, we determined pathological differences in the 5xFAD mouse visual system allowing us to identify Aβ-specific effects to microglia in visual regions of the brain. Examining how regions in the image-forming pathway for vision (dLGN, V1) and non-image-forming regions (SC, SCN) respond to Aβ plaque formation is critical to our understanding of how AD affects vision as deficits in vision manifest early in AD patients, how Aβ plaques are removed from the brain during AD, and how microglia contribute to disease mitigation or progression.

## 2. MATERIALS AND METHODS

### 2.1 Animals

Mice were housed on a 12 hr light/dark cycle at the University of Nebraska Medical Center with professional veterinary support through the Department of Comparative Medicine and access to food and water *ad libitum*. 5xFAD mice [B6.Cg-Tg(APPSwFlLon,PSEN1*M146L*L286V)6799Vas/Mmjax (5XFAD, The Jackson Laboratory #034848-JAX, RRID:MMRRC_034848-JAX] were purchased from Jackson Laboratories. These mice were on a C57BL/6J background and did not contain the retinal degeneration allele Pde6b^rd1^. A recent study has shown that parental origin of the transgene plays a role in Aβ plaque burden (Sasmita et al., 2025). As our colony may have received the transgene from either parent, some variability in our study population may be due to parental origin of the transgene. C57BL/6J mice (C57, The Jackson Laboratory #:000664, RRID:IMSR_JAX:000664)] were used as age-matched controls. Both male and female mice were used to account for sex differences. All animal protocols were approved by the Institutional Animal Care and Use Committee at the University of Nebraska Medical Center. Mice were sacrificed via CO_2_ inhalation followed by cervical dislocation as the secondary method of euthanasia.

### 2.2 Histology and immunohistochemistry

Brains were fixed by submersion in 4% paraformaldehyde for 4 hrs, cryo-protected in 30% sucrose in 0.01 M PBS, and embedded in 3% agar. Using a Leica VT1000S vibratome, 50-μm-thick sections containing dLGN, V1, SC, or SCN were created, mounted on Superfrost Plus slides (Fisher), and stored at −20 °C prior to staining.

#### 2.2.1 Thioflavin S staining

To stain sections for Aβ plaques, slides were washed 1 x 1 min in 70% ethanol, 1 x 1 min in 80% ethanol, 1 x 15 min in a filtered (0.2 μm) thioflavin S solution, 1 x 1 min in 80% ethanol, 1 x 1 min in 70% ethanol, and 2 x 1 min in dH_2_O. Coverslips were mounted using Vectashield Hardset mounting medium and stored at 4 °C until imaged.

#### 2.2.2 Iba1 and CD68 immunostaining

We used two markers to examine different aspects of the role of microglia in the brain. Ionized calcium binding adaptor molecule 1 (Iba1) was used to label microglia for measurements of morphology while cluster of differentiation 68 (CD68) was used as a marker of phagocytic activity. Sections were washed 1 x 1 min in 0.01 M PBS followed by a 1 hr incubation in 150 μL of blocking solution (0.01 M PBS, 20% Triton X-100, donkey serum, goat serum) in a humidified chamber at room temperature. Tissue sections were then incubated in 150 μL primary antibody (1:500 rabbit anti-Iba1, Wako CAT# 019-19741, RRID:AB_839504; 1:1000 rat anti-CD68 Abcam CAT #ab53444, RRID:AB_869007) in a humidified chamber at 4 °C for 3 nights. Slides were washed 6 x 10 min in 0.01 M PBS followed by another 1 hr incubation in 150 μL of blocking solution. After blocking, slides were incubated for 2 hrs in 1:200 secondary antibody (goat anti-rat 488, Thermo Fisher Scientific CAT #A11006, RRID:AB_2534074; donkey anti-rabbit 568, Thermo Fisher Scientific CAT # A10042, RRID:AB_2534017), then washed 3 x 10 min in 0.01 M PBS and 1 x 1 min in dH_2_O. Tissue was coverslipped with Vectashield hardset mounting medium and stored at 4 °C until imaged. Tissue used for skeleton analysis of 6, 9, and 12 mo dLGN tissue was not blocked prior to incubation in secondary antibody.

### 2.3 Imaging and analysis

#### 2.3.1 Thioflavin S imaging and analysis

Images of relevant tissue regions were captured using a fixed-stage upright microscope (Olympus BX51-WI). To quantify Aβ plaque density in dLGN, SC, and SCN, ImageJ was used to measure the area of each brain region, and the Cell Counter plug-in was used to count Aβ plaques. For V1, analysis of Aβ density of a 1280 x 1024 pixel (65 pixels / 50 μm) image was analyzed, and the Cell Counter plug-in was used to count Aβ plaques. Aβ plaques were also analyzed as a percentage of the depth of V1.

#### 2.3.2 Iba1 and CD68 imaging and analysis

For Iba1- and CD68-stained tissue sections, a 370 μm x 370 μm (1024 x 1024 pixel) field of view with 1 μm z-spacing and 4 frames per plane for each brain region was imaged using a 2-photon microscope with a Ti:sapphire laser tuned to 800 nm.

##### 2.3.2.1 Iba1 skeleton analysis protocol

After image acquisition, 4 frames per plane were averaged resulting in a stack from which 40 optical sections of 1 μm thickness, giving a 40 μm z-depth, were kept for analysis. Images were converted to 8-bit grayscale, contrast was adjusted, image was sharpened and despeckled, then a fixed manual threshold was applied corresponding to 22% of the maximum intensity. Using ImageJ, the microglia were skeletonized to find endpoints and branch length via the Analyze Skeleton (2D/3D) function. The Cell Counter plugin was then used to count microglia to calculate endpoints and branch length per microglia. Our lab has previously used this method of analysis for microglia in the dLGN of other mouse models following previously published methods (Bhandari et al., 2022; Thompson et al., 2025; Morrison et al., 2017; Young and Morrison, 2018).

##### 2.3.2.2 Iba1/CD68 colocalization analysis protocol

Channels were thresholded separately and then merged. Pixel colocalization was measured as the ratio of colocalized pixels (cyan) to colocalized pixels plus Iba+ only (blue) pixels in a single plane. To account for random colocalization, a single channel was rotated 90°, and the analysis was repeated.

### 2.4 Statistical analyses

GraphPad Prism 10 software was used for statistical analyses. Data were tested for normality using a D’Agostino and Pearson test. When normally distributed, a 2way ANOVA with multiple comparisons, an unpaired t-test, or a nested t-test were used. When not normally distributed, a Kruskal-Wallis test was used. Figure legends contain information on statistical tests used, p-values, and sample sizes.

## 3. RESULTS

### 3.1 A**β** Plaque Density by Visual Brain Region

We first sought to examine the extent to which Aβ plaque pathology is present in regions of the brain responsible for distinct visual tasks (Tsui et al., 2022). We quantified Aβ plaque density in each visual brain region by staining 50-μm-thick tissue sections containing the region of interest using thioflavin S (Fig. 1.A). The area of each brain structure was measured, with the exception of V1 where only a portion was analyzed, and Aβ plaques were counted manually within that area to determine Aβ plaque density. C57 mice presented with no Aβ plaques in all regions examined. Aβ plaque density was higher in the dLGN compared to C57 mice as well as compared to other visual brain regions. There was a significantly higher density of Aβ plaques in V1 compared to the SC and SCN, however, the density of Aβ plaques in V1 was much less than in the dLGN. We saw no Aβ plaques in the SCN which aligns with previously reported work in this model (Nam et al., 2022). 5xFAD SC tissue showed minimal Aβ plaque deposition, with only a few sections containing one or two Aβ plaques (Fig. 1.B). Previously, we showed that there is a significant density of Aβ plaques present in the dLGN from 6 mo onward with no change in count density up to 12 mo (McCool et al., 2025). Since we saw no change in number of Aβ plaques, we decided to look at percent area of Aβ plaque coverage to determine whether plaques were changing in size. Analyzing the dLGN at 6, 9, and 12 mo, we found a significant increase in percent area of the dLGN covered in Aβ plaques at all time points compared to age-matched controls (Fig. 1.D). We also saw an increase in coverage from 6 mo to 12 mo and from 9 mo to 12 mo, but not between 6 mo and 9 mo. For comparison to a disease-relevant region, we analyzed percent area of hippocampus covered in Aβ plaques finding similar results. These data point to a spatiotemporal path of Aβ plaque deposition in 5xFAD visual brain regions with minimal Aβ plaque deposition in the SC and SCN but significant Aβ plaque burden in V1 and increasing Aβ plaque size with age in the dLGN.

**Fig. 1.**
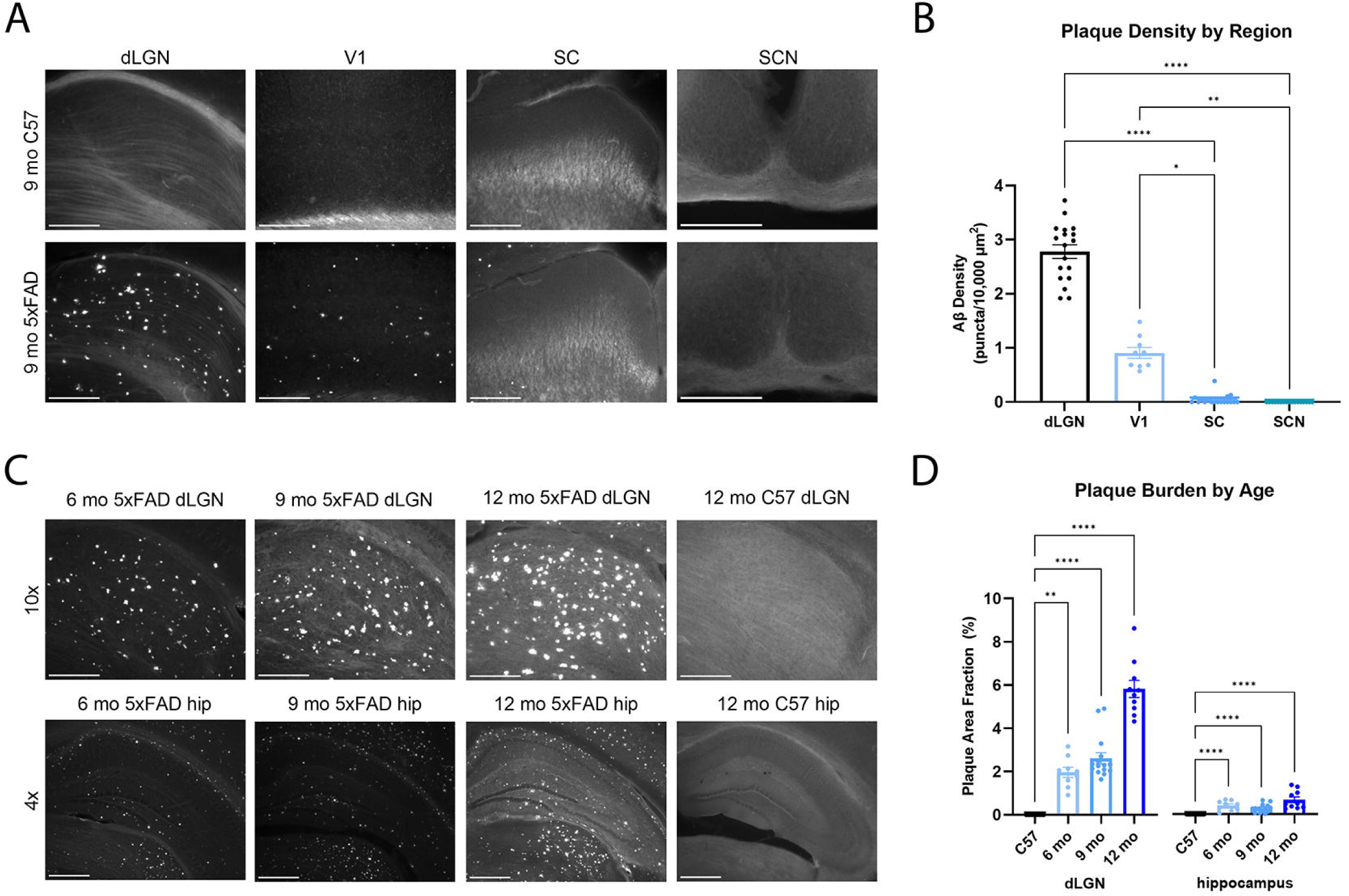
Analysis of Aβ plaque density reveals differential Aβ plaque deposition in visual regions of the brain. **(A)** Example images of 9 mo C57 and 5xFAD dLGN, V1, SC and SCN stained for Aβ using thioflavin S to show Aβ plaque density in visual brain regions. **(B)** Aβ plaque density by region reveals significant Aβ plaque formation in 5xFAD dLGN and V1 with minimal to no Aβ plaque formation in SC and SCN as analyzed by one-way ANOVA. **(C)** Example images of 6, 9, and 12 mo 5xFAD dLGN and hippocampus and 12 mo C57 dLGN and hippocampus stained for Aβ plaques using thioflavin S to show Aβ plaque coverage over time. **(D)** Aβ plaque coverage over time reveals a significant Aβ plaque burden in 5xFAD dLGN and hippocampus at all time points and significant increase from 6 mo to 12mo and 9 mo to 12 mo. Analysis was performed on both left and right hemispheres. Sample size described as number of mice: n = 9 5xFAD for dLGN, V1, and SC; n = 6 5xFAD for SCN. Scale bar = 250 μm. For (B), p < 0.0001 for all statistical measures shown. For (C), p < 0.0001 for all statistical measures shown for dLGN. For hippocampus, p < 0.0001 for all statistical measures shown between C57 and 5xFAD mice and between 9 mo and 12 mo 5xFAD mice, and p = 0.0054 for 6 mo and 12 mo 5xFAD comparison.

**Fig. 2.**
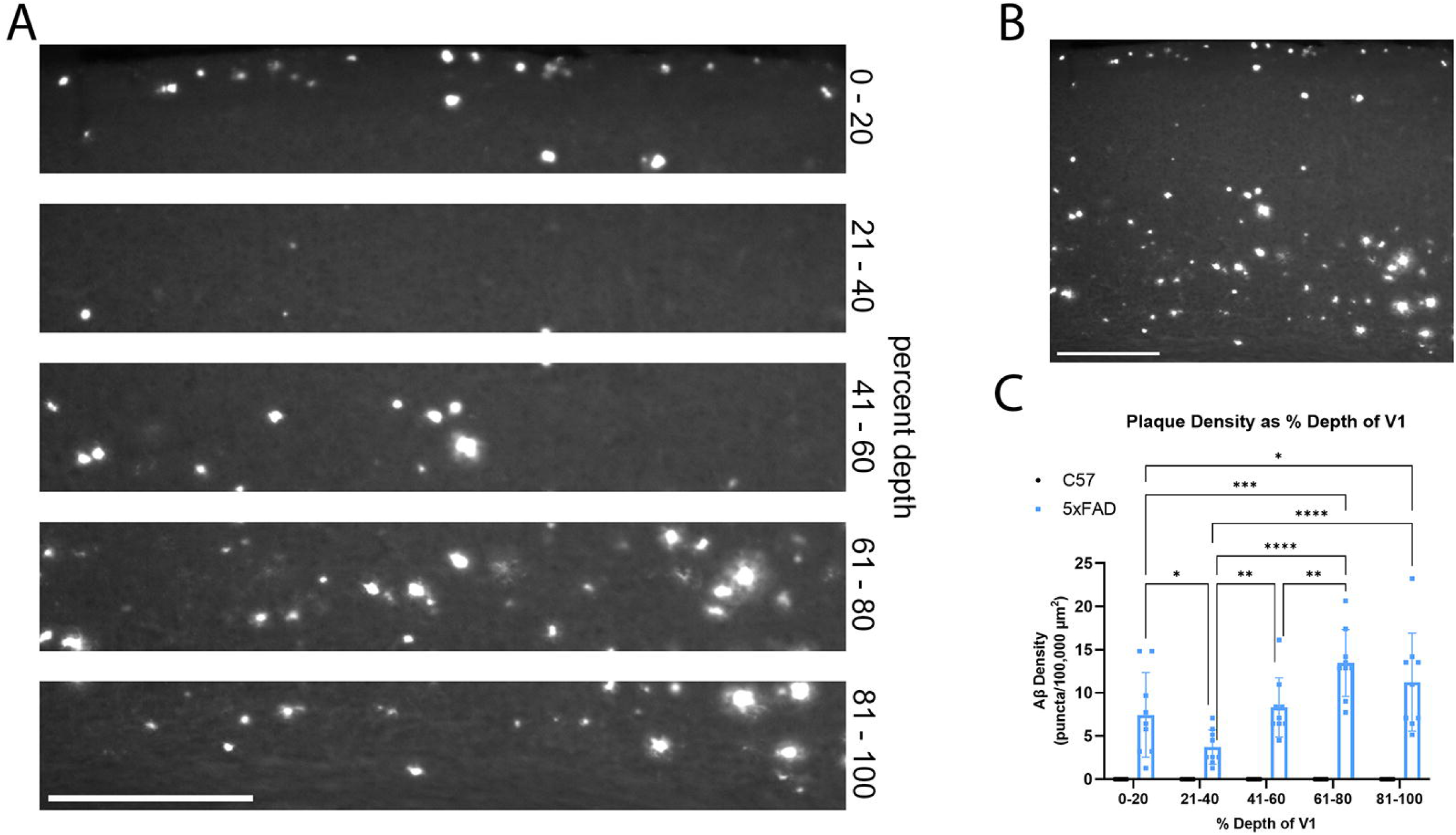
Analysis of Aβ plaque density in V1 reveals varied Aβ plaque density by depth. **(A)** Example image of V1 split into 20% increments for analysis of Aβ plaque density by percent depth of V1. **(B)** Example from (A) showing full image. **(C)** Quantification of Aβ plaque density as a percent depth of V1 shows significant changes in Aβ plaque density depending on depth as analyzed by 2way ANOVA with multiple comparisons. Sample size described as number of mice: n = 8 C57 and n = 9 5xFAD. For genotype comparisons, p < 0.0001 (0 – 20, 41 – 60, 61 – 80, and 81 – 100%) and p = 0.0139 (21 – 40%). For depth comparisons, p < 0.0001 (21 – 40 vs. 41 – 60% and 81 – 100%), p = 0.0001 (0 – 20% vs. 61 – 80%), p = 0.0057 (21 – 40% vs. 41 – 60%), p = 0.0014 (41 – 60 vs. 61 – 80%), p = 0.0378 (0 – 20% vs. 21 – 40%), and p = 0.327 (0 – 20% vs. 81 – 100%).

### 3.2 V1 Shows Differential A**β** Plaque Density Based on Cortical Depth

Due to the stratified nature of V1, we were interested in whether Aβ plaque density was altered based on cortical depth. To accomplish this, each image of V1 was divided into 5 sections from the pial surface to the ventral side of layer 6 with each section representing 20% of the image in order to analyze Aβ plaque density as a function of depth in V1. In each depth bin of V1, we found that Aβ plaque density was significantly higher at all depths in 5xFAD V1 compared to age-matched controls (Senzai et al., 2019). Additionally, we found a differential density of Aβ plaques depending on cortical depth within 5xFAD samples, with Aβ plaque density being significantly lower at the 21 – 40% depth, which corresponds approximately to layer 2/3, compared to other layers. We also found a significantly higher Aβ plaque density at 61 – 80% depth compared to the first 60% of V1 but similar density compared to 81 – 100% depth. 61 – 100% depth corresponds to layers 5 and 6 indicating they have a significantly higher Aβ plaque density compared to layers 1 – 4. From 81 – 100% depth, we found a significantly higher Aβ plaque density compared to the first 40% of tissue but comparable to 61 – 80% depth. For 41 – 60% depth, corresponding to layer 4, we found comparable Aβ plaque density to the first and last 20% but significantly distinct from neighboring depths – more than 21 – 40% but less than 61 – 80%. Lastly, the first 20% of the tissue is significantly impacted by Aβ plaques compared to 21 – 40% while being less impacted compared to 61 – 100% depth. These results show that distinct layers of the primary visual cortex are differentially impacted by AD-like pathology.

### 3.3 Microglia Cluster Around A**β** Plaques in Specific Visual Brain Regions

Microglia are implicated in AD pathology, even showing Aβ-specific profiles (Gerrits et al., 2021). To determine the role of microglia in visual brain regions of the 5xFAD mouse, we first tested whether microglia localize with Aβ plaques in the regions displaying significant Aβ plaque deposition. The 5xFAD dLGN and V1 were costained for microglia and Aβ plaques using Iba1 and thioflavin S, respectively. Images of each fluorophore were taken separately and merged. In the dLGN and V1, regions with significant Aβ plaque density, we found that microglia clustered around thioflavin S-labeled Aβ plaques (Fig. 3.A). The SC and SCN were not analyzed for localization of microglia and Aβ plaques due to the relatively insignificant density of Aβ plaques present in these regions. Having previously reported significant Aβ plaque deposition in the 5xFAD dLGN at 6, 9, and 12 mo (McCool et al., 2025), and in this study showing significant Aβ plaque deposition in V1, we assessed whether microglia density was altered at these ages and in the visual brain regions of interest. We found a significantly higher density of microglia in the 5xFAD dLGN at all time points compared to age-matched controls (Fig. 3.B) as well as in V1 while the SC and SCN showed no change in microglial density (Fig. 3.C). In dLGN, which we were able to analyze at 6-, 9-, and 12-mo timepoints, microglial density increased over time, with higher density at both 9 and 12 mo compared to 6 mo. These observations suggest a relationship between microglia and Aβ plaques in the 5xFAD mouse, exemplifying the intertwining nature of microglia and Aβ plaque pathology in AD.

**Fig. 3.**
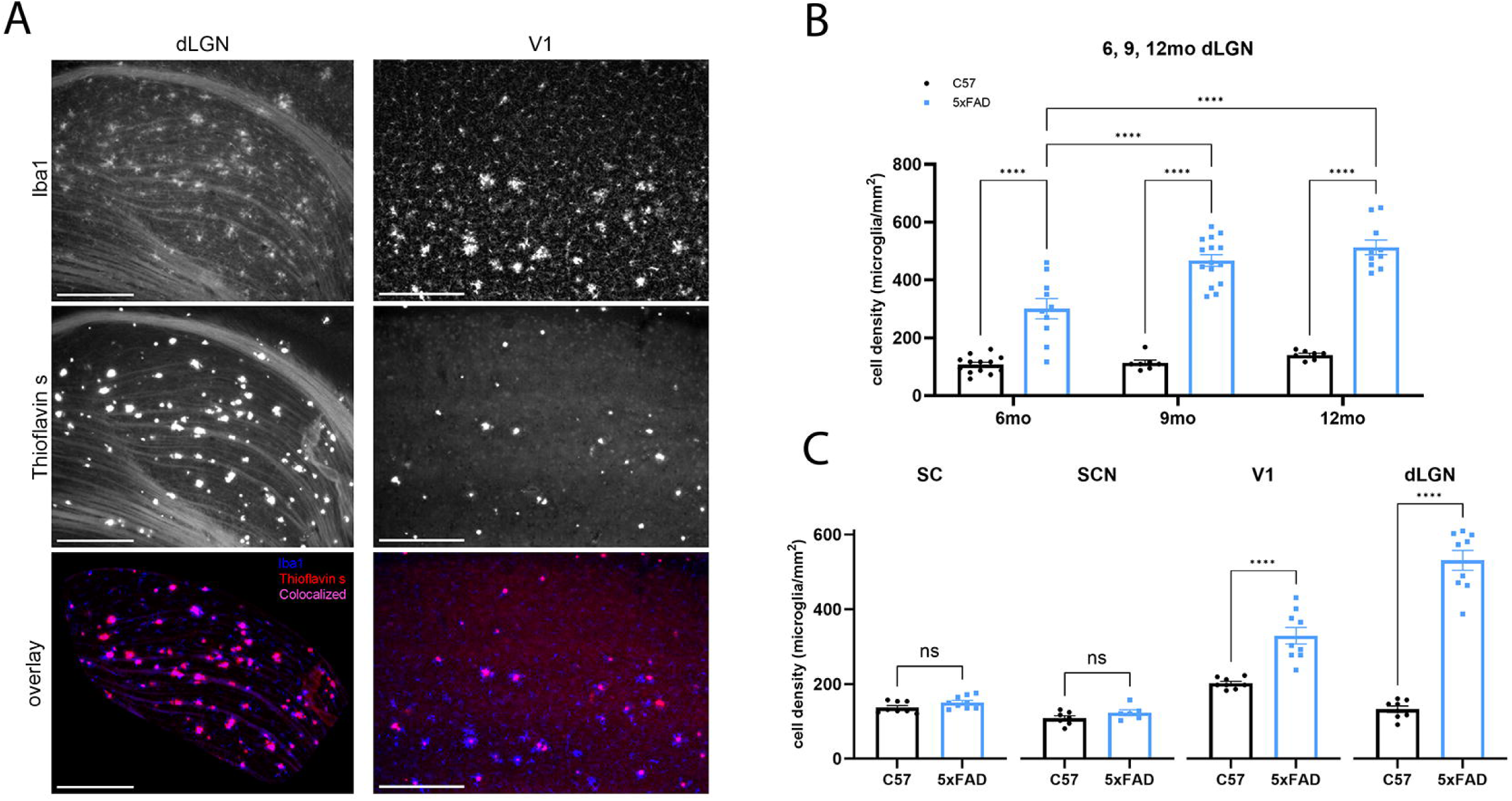
Microglia cluster around Aβ plaques in the thalamocortical pathway of 5xFAD mice in addition to experiencing increased microglia cell density. **(A)** Example images of dLGN and V1 stained for Iba1 and Aβ plaques to show relative localization of microglia and Aβ plaques. **(B)** Microglia cell counts in 6, 9, and 12 mo dLGN of C57 and 5xFAD mice are significantly increased at all time points in the 5xFAD mice compared to controls as well as between time points as analyzed via 2way ANOVA with multiple comparisons. **(C)** Microglia cell counts in 9 mo dLGN, V1, SC, and SCN of C57 and 5xFAD mice are significantly increased in the 5xFAD dLGN and V1 as analyzed via unpaired t-test. For (B), one image per brain was analyzed. For (C), two images per brain with one image of each bilateral region were analyzed. Sample size described as number of mice: n = 13 (6 mo C57) 10 (9 mo C57) 7 (12 mo C57) 10 (6 mo 5xFAD) 15 (9 mo 5xFAD) 10 (12 mo 5xFAD); n = 8 C57 and 9 5xFAD for 9 mo dLGN, V1, and SC; n = 7 C57 and 6 5xFAD for 9 mo SCN. Scale bar = 250 μm. Cell density p-values: For (B), p < 0.0001 for all statistical measures shown. For (C), p < 0.0001 for dLGN and V1; p = 0.1829 for SCN; p = 0.0963 for SC.

### 3.4 Skeleton Analysis Reveals Shifts in Microglia Morphology in Plaque-infested Visual Brain Regions

After establishing that microglia cluster around Aβ plaques in affected regions, we aimed to examine microglial morphology, a meaningful indicator of their activity, in our regions of interest. Using Iba1 as a marker for microglial morphology, we performed a skeleton analysis to assess both the branch length and endpoints per microglia (Fig. 4.A) (Morrison et al., 2017; Young and Morrison, 2018; Bhandari et al., 2022; Thompson et al., 2025). We examined the dLGN at three time points to determine whether there are age-associated changes in microglial morphology associated with higher Aβ plaque burden. We found a significantly lower branch length and number of endpoints per microglia at all three time points in the 5xFAD dLGN compared to age-matched controls but did not detect a significant difference across time points within the 5xFAD samples (Fig. 4.B). There was an effect of age on the number of endpoints and branch length in the C57 samples, which parallels similar age-dependent changes we have observed in the dLGN of otherwise healthy mice in a study of glaucoma (Thompson et al., 2025). We repeated this skeleton analysis in all four brain regions at the 9-mo time point to examine regional differences in microglial morphology (Fig. 5.A). Skeleton analysis of the dLGN, V1, SC, and SCN revealed significantly lower branch length and endpoints per microglia in the dLGN and V1, regions that also had high Aβ plaque density, of the 5xFAD brains compared to C57 control tissue. The two regions with low to no Aβ plaque deposition, SC and SCN, showed no significant difference in either morphological parameter compared to controls (Fig. 5.B). Overall, these results demonstrate the dynamic nature of microglia particularly in response to Aβ pathology, in their environment.

**Fig. 4.**
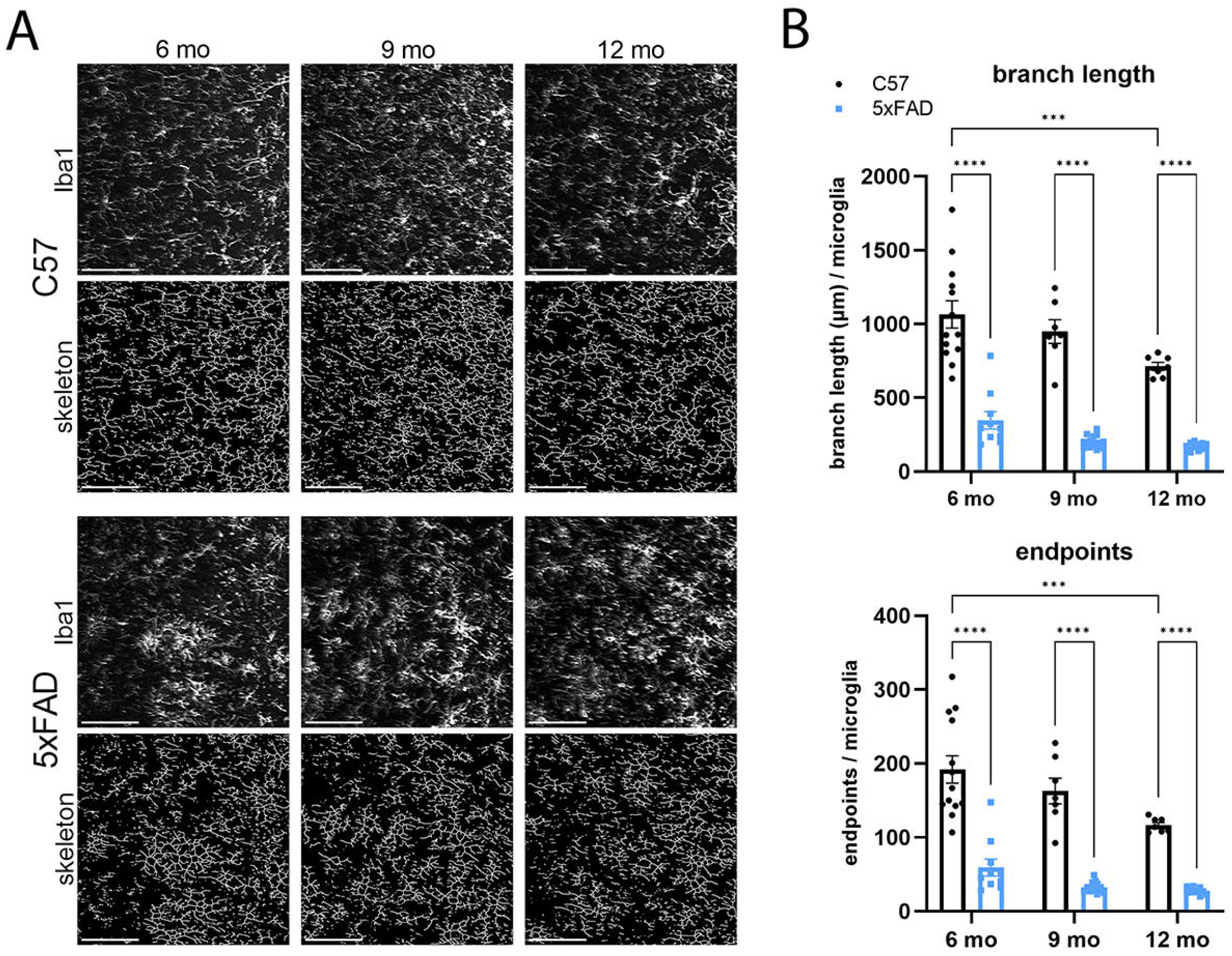
Microglia in the dLGN display amoeboid morphology at all timepoints. **(A)** Example images of visual brain regions stained for Iba1 with skeletonized image in 6, 9, and 12 mo C57 and 5xFAD mice. **(B)** Quantification of branch length and endpoints per microglia in 6, 9, and 12 mo C57 and 5xFAD mice shows significantly decreased parameters in the dLGN at all timepoints as analyzed via 2way ANOVA with multiple comparisons. Two images per brain, with one image of each bilateral region, were analyzed. Sample size described as number of mice: n = 13 (6 mo C57) 7 (9 mo C57) 7 (12 mo C57) 10 (6 mo 5xFAD) 15 (9 mo 5xFAD) 10 (12 mo 5xFAD). Scale bar = 100 μm. For time point comparisons: p < 0.0001. For genotype comparisons: p = 0.0002 (6 vs 12 mo C57 endpoints); p = 0.0006 (6 vs 12 mo C57 branch length).

**Fig. 5.**
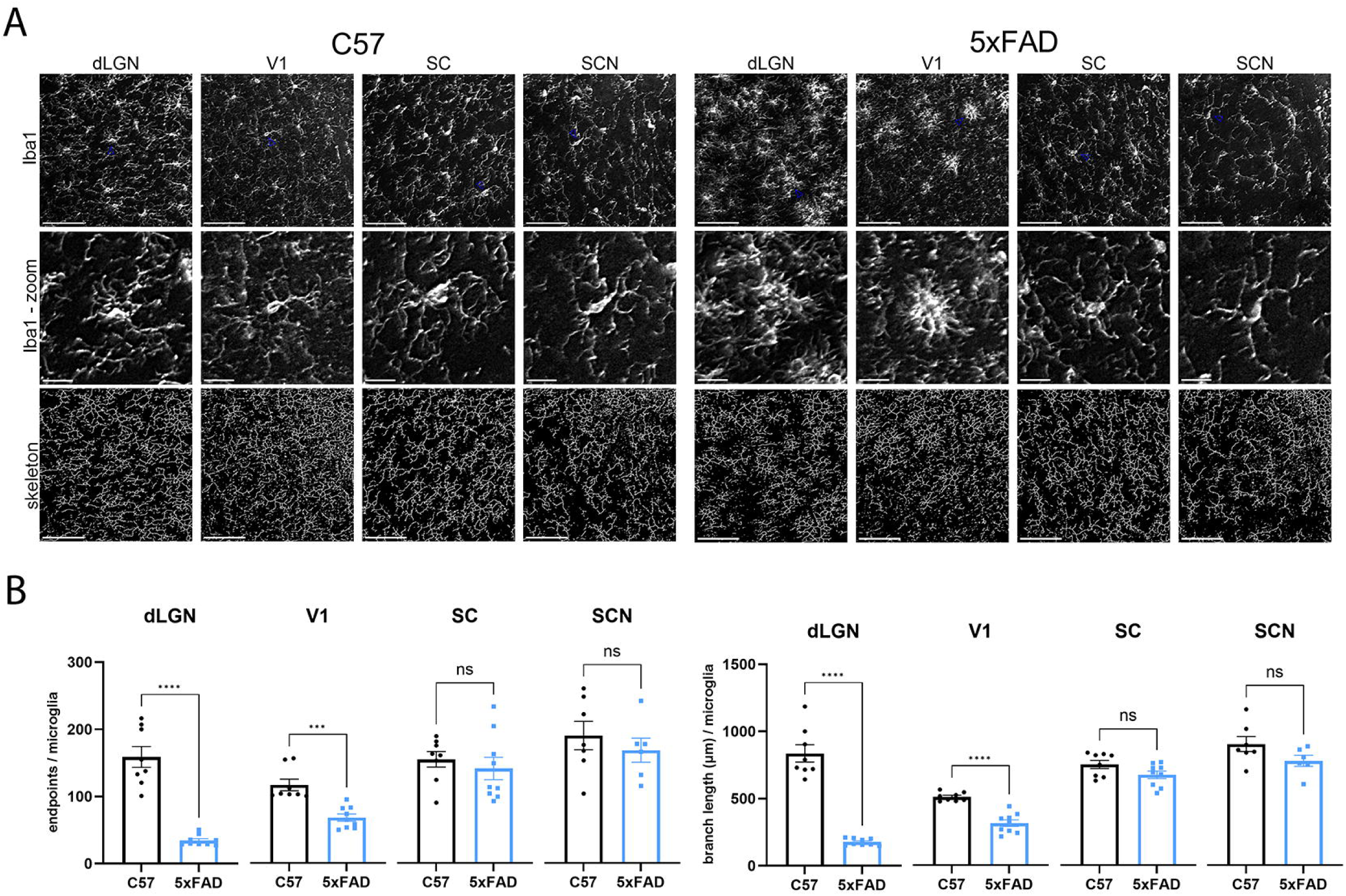
Microglia in the dLGN and V1 display amoeboid morphology. **(A)** Example images of visual brain regions stained for Iba1 with zoomed in image of a single microglia in each region as well as a skeletonized image in 9 mo C57 and 5xFAD mice. **(B)** Quantification of branch length and endpoints per microglia in C57 and 5xFAD mice shows a significant decrease in the dLGN and V1 but not in SC or SCN as analyzed via nested t-test (data represented as t-test). Two images per brain, with one image of each bilateral region, were analyzed. Sample size described as number of mice: n = 8 C57 and 9 5xFAD for dLGN, V1, and SC; n = 7 C57 and 6 5xFAD for SCN. Scale bar = 100 μm. Branch length: p < 0.0001 for dLGN and V1; p = 0.0828 for SC; p = 0.1074 for SCN. Endpoints: p < 0.0001 for dLGN; p = 0.0002 for V1; p = 0.5261 for SC; p = 0.4577 for SCN.

### 3.5 Colocalization Analysis Reveals Phagocytic Microglia in Plaque-infested Visual Brain Regions

The highly plastic nature of microglia allows their morphology to be used as a window into their functional state. Microglial reactivity is often associated with increased phagocytic activity. Therefore, to test how the morphology is reflected in functional phenotype, we analyzed phagocytic profiles by testing for colocalization of Iba1-labeled microglia with CD68, a lysosomal protein indicative of increased phagocytic activity. We first costained the dLGN at 6, 9, and 12 mo to determine age-associated changes in the microglial response to Aβ plaque formation (Fig. 6.A). We found a significant colocalization of Iba1 and CD68 in 9- and 12-mo 5xFAD dLGN, but not in 6-mo 5xFAD dLGN, compared to age-matched controls (Fig. 6.B). We performed an analysis to control for random colocalization by rotating one channel 90° after which we saw the colocalization significantly decreased at all time points and in all analyzed brain regions indicating random colocalization is minimal (Fig. 6.C). In order to determine regional differences in microglial activity, we repeated the analysis at 9 mo in all four visual brain regions (Fig. 7.A). The dLGN and V1 both showed significantly higher colocalization of Iba1 and CD68 in the 5xFAD tissue relative to C57 controls, but this was not the case in the SC or SCN (Fig. 7.B). This follows the previous pattern seen in the skeleton analysis where we identified decreased branch length and endpoints in the dLGN and V1 but not in the SC or SCN. These findings support the idea that microglia are in a phagocytic state specifically in regions of the brain affected by AD-like pathology.

**Fig. 6.**
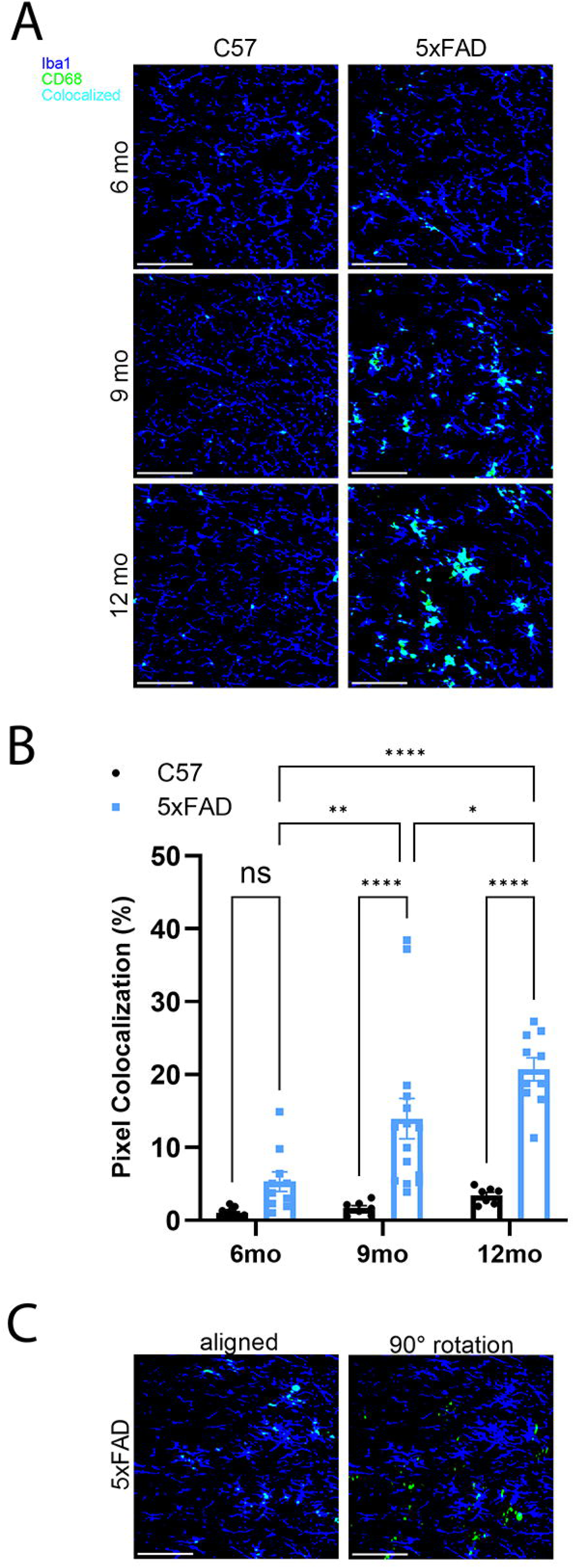
Increased phagocytic activity is seen in the dLGN of 5xFAD mice as they age. **(A)** Example images of 6, 9, and 12mo C57 and 5xFAD dLGN stained for Iba1 and CD68 show microglial phagocytic activity. **(B)** Colocalization of Iba1 and CD68 is significantly different between 9 mo and 12 mo C57 and 5xFAD mice and between 5xFAD 6, 9, and 12 mo time points analyzed via 2way ANOVA with multiple comparisons. **(C)** Example images showing Iba1 and CD68 colocalization (left) and disappearance of colocalization when one channel is rotated 90°. Sample size described as number of mice: n = 13 (6 mo C57) 10 (6 mo 5xFAD) 7 (9 mo C57) 15 (9 mo 5xFAD) 7 (12 mo C57) 10 (12 mo 5xFAD). Scale bar = 100 μm. For time point comparisons: p = ns (6 mo); p < 0.0001 (9 mo); p < 0.0001 (12 mo). For genotype comparisons: p = ns (C57); p = 0.0026 (6 vs 9 mo 5xFAD); p < 0.001 (6 vs 12 mo 5xFAD); p = 0.0231 (9 vs 12 mo 5xFAD).

**Fig. 7.**
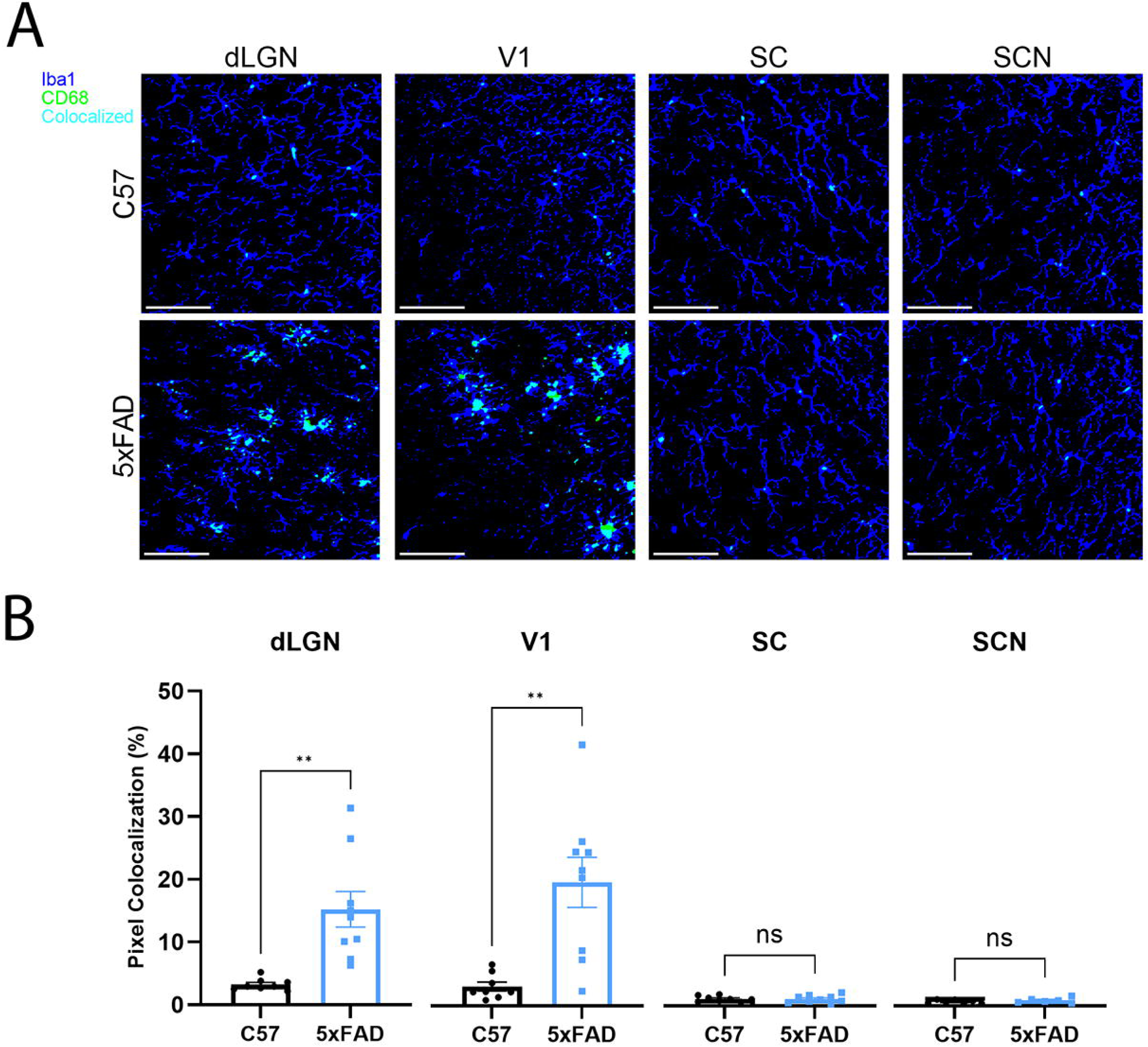
Microglia in visual brain regions with significant Aβ plaque deposition show increased phagocytic activity. **(A)** Example images of C57 and 5xFAD dLGN, V1, SC, and SCN stained for Iba1 and CD68 show microglial phagocytic activity. **(B)** Colocalization of Iba1 and CD68 is significantly increased in 5xFAD dLGN and V1 compared to C57 mice as analyzed via a nested t-test. Two images per brain with one image of each bilateral region were analyzed with results averaged for analysis. Sample size described as number of mice: n = 8 C57 and 9 5xFAD for dLGN, V1, and SC; n = 7 C57 and 6 5xFAD for SCN. Scale bar = 100 μm. p = 0.0013 for dLGN; p = 0.0015 for V1; p = 0.8945 for SC; p = 0.3181 for SCN.

## 4. DISCUSSION

Visual dysfunction is an identified pathological occurrence in AD patients. Many of these visual deficits occur early in disease progression similar to Aβ pathology which is also one of the earliest events in AD pathogenesis. AD patients experience decreases in contrast sensitivity, color vision, visual acuity, and visual field which are visual functions associated with the image-forming pathway including the lateral geniculate nucleus and V1 (Javaid et al., 2016). AD patients also experience circadian rhythm disruptions, oculomotor dysfunction, and impaired multisensory integration which are tasks associated with non-image-forming regions, specifically the SC and SCN (Javaid et al., 2016; Nassan et al., 2021; Mahoney et al., 2025). It is unclear whether visual dysfunction in AD depends on both Aβ plaques and NFTs as both pathologies have been identified in human retina and visual brain regions as well as in the retina and visual brain regions of many AD mouse models. There are also conflicting reports on whether microglia, the immune cells responsible for protecting neurons, contribute to the progression or elimination of amyloid pathology. Some reports show microglia degrade Aβ via digestive exophagy, limit Aβ plaque size, or increase phagocytosis via modulation of neuropeptides or during glymphatic system impairment and therefore work synergistically to prevent Aβ plaque formation, results which indicate that they are beneficial to Aβ plaque removal (Fleisher-Berkovich et al., 2010; Zhao et al., 2017; Feng et al., 2020; Jacquet et al., 2024). Conversely, some research shows depletion of microglia leads to reduced Aβ plaques or that Aβ-activated microglia induce the spread of AD pathology indicating their detrimental role in AD pathogenesis (Spangenberg et al., 2019; Baligács et al., 2024; Lee et al., 2024). In between these two opposing theories, others demonstrate that the depletion of microglia has no effect on amyloid pathology (Spangenberg et al., 2016; Gratuze et al., 2021). Although their role in Aβ pathology is unclear, they have been shown to contribute to pathological synaptic pruning in AD and frontotemporal dementia acting as drivers of neurodegeneration (Hong et al., 2016; Lui et al., 2016). Therefore, microglia play an unknown role in the AD brain and may have a bidirectional impact.

For that reason, we aimed to identify morphological and behavioral changes in microglia in visual brain regions in a model of Aβ pathology. In this study, we utilized immunohistochemistry and 2-photon imaging to approach the question of whether brain regions responsible for visual tasks and computations experience differential Aβ plaque deposition in AD and how microglia are responding to this AD-like pathology. The major finding of this study is that visual brain regions in the pathway for conscious vision, the dLGN and V1, have significant Aβ plaque deposition that coincides with microglial polarization, particularly to an amoeboid state, and these microglia exhibit phagocytic activity in an age-dependent manner. However, visual regions responsible for non-image forming vision, the SC and SCN, show minimal Aβ plaque formation with ramified, non-phagocytic microglia. Our findings demonstrate specific, disease-associated features of microglia within visual brain regions experiencing, or not experiencing, AD-like pathology.

Our findings show significant Aβ plaque density in several brain regions important for vision-dependent tasks. This includes the dLGN at 6, 9, and 12 mo as well as V1 at 9 mo. In contrast, the SC and SCN show either minimal or no Aβ plaque formation, respectively. These findings are consistent with prior research in the 5xFAD mouse model which shows Aβ plaque formation in the dLGN and V1 but no Aβ plaque formation in the hypothalamus where the SCN is located (Tsui et al., 2022; Nam et al., 2022). However, these studies found no Aβ plaques present in the SC while we found a few Aβ plaques, although insignificant. In humans, Aβ plaques have been found in the SC, LGN, and occipital cortices, including V1, with some Aβ plaques present in the SCN, although no mature neuritic plaques were present (Brilliant et al., 1992; Stopa et al., 1999; Erskine et al., 2015; Ikonomovic et al., 2016; Hwang et al., 2021). Our data are intriguing as they show significant AD-like pathology in the pathway for conscious vision but minimal pathology in visual regions used in non-conscious visual processing. Due to the progressive nature of AD, we also aimed to look at Aβ pathology in the dLGN, the thalamic relay for conscious vision, at three ages to resolve the pathological progression in this region over time. Given that we previously saw no change in Aβ plaque count in the dLGN from 6 mo to 12 mo, we would assume that the regions currently devoid of Aβ plaques, the SC and SCN, would present with Aβ plaques by the 9-month time point if they were to develop any (McCool et al., 2025). As they show either insignificant or no Aβ plaque density by 9 mo, we suspect they may not develop Aβ plaques. This region-specific amyloid pathology could be a result of the genetics of the mouse model. The promoter used for this model, Thy1, is expressed in neurons, and therefore directs expression widely throughout the brain, although it is not a uniform expression (Moechars et al., 1999; Feng et al., 2000; Oakley et al., 2006). The Thy1 promoter is regularly used to investigate neuronal activity in mouse lines, and despite the fact that expression is not directly linked to being regulated by neuronal activity, these studies suggest that more active brain regions will have higher expression of the transgenes (Chen et al., 2012; Cichon et al., 2020). Additionally, it has been shown that transgene inheritance in this model plays a role in Aβ plaque burden which suggests it could play a role in Aβ plaque patterning, both temporally and spatially, as well (Gregg et al., 2010; Sasmita et al., 2025). Therefore, the pathway-specific divergence in amyloid pathology might be attributable to genetic characteristics of the model. Furthermore, the differential Aβ plaque deposition does not seem to be a fundamental feature of AD: first, the Aβ plaque deposition seen in this model does not directly follow that of human AD and, second, Aβ plaques have been identified in all four of these visual brain regions in AD patients (Palmqvist et al., 2017; Oakley et al., 2006; Brilliant et al., 1992; Stopa et al., 1999; Erskine et al., 2015; Ikonomovic et al., 2016; Hwang et al., 2021). When exposed to Aβ, the SC experiences retinal ganglion cell axon terminal degeneration and gliosis (Simons et al., 2021). Given this fact, Aβ mediates activation of microglia in this region as well, although not shown in the 5xFAD model. Alternatively, thalamocortical and corticothalamic transport between the dLGN and V1 are potential factors contributing to a larger Aβ plaque burden in the conscious visual pathway. The dLGN projects to layer 4 of V1 while layer 6 of V1 projects back to the dLGN. Reciprocal transport of Aβ between dLGN and V1 may contribute to the larger Aβ plaque burden in these regions as Aβ has been shown to spread through synaptically-connected regions (Song et al., 2014; Pignataro et al., 2017). Interestingly, we also see the largest Aβ plaque density at 61 – 100 % depth which incorporates portions of cortical layers 5 and 6 and is consistent with initial reports of Aβ plaque deposition in this model (Oakley et al., 2006). Thus, the cortical layers of V1 most intensely affected by AD-like pathology are also the layers contributing input to dLGN. Overall, these data point to significant AD-like pathology in the geniculo-cortical pathway for conscious, image-forming vision but not in regions responsible for non-image-forming aspects of vision in the 5xFAD mouse.

The observed trends indicate that cortical layer 2/3 is the least affected by Aβ pathology in the 5xFAD mouse model with deeper layers being significantly more affected. Layer 5 of the cortex has been shown to have significant neuron loss as well as the highest levels of both intraneuronal Aβ accumulation and Aβ deposition in one of the original 5xFAD lines (Tg6799) (Oakley et al., 2006). This may be why we see a significantly higher density of Aβ plaques at 61-80% depth, correlated with layer 5 depth, of V1. Developmental timing may also play a role as early-born neurons such as those in layers 5 and 6 have a longer duration of Thy1 promoter activity as Thy1 is a marker of mature neurons (Gregg et al., 2010). As specific layers of V1 are differentially impacted by AD-like pathology, this suggests that specific visual behavioral outputs may be differentially affected in these mice as well.

In an effort to determine whether microglia are attempting to remove Aβ plaques, we assessed whether microglia colocalize with Aβ plaques, indicative of neuritic Aβ plaque formation (Tsering et al., 2025). Our findings show that microglia cluster around Aβ plaques in the regions with significant Aβ plaque burden. Within these same regions, we see a significant increase in microglia counts. We also see significantly increased microglia counts over time in the dLGN. Previously, we saw significant Aβ plaque burden in the dLGN at all timepoints, but no change in Aβ plaque density by count over time (McCool et al., 2025). However, we show that although Aβ plaque density by count does not increase over time, Aβ plaque coverage of the dLGN does increase over time. Microglial density and CD68 labeling also show a lag relative to Aβ plaque density as both these parameters increase over time similar to Aβ plaque size. Thus, increasing Aβ plaque size may be a factor in microglia presence in dLGN, contributing to increased microglial count over time. With V1, we see microglia clustering around plaques and increased microglia counts compared to other regions at 9 mo indicating significant Aβ plaque presence. These data are indicative of Aβ plaque deposition driving microglial recruitment. A disease-relevant region, the hippocampus, shows similar pathology in terms of Aβ plaque deposition and microglial clustering, and therefore likely displays similar morphological and phagocytic changes to the visual brain regions that also display significant pathology. Newly published data show no change in hippocampal Iba1, CD68, microglial density, or microglial morphological parameters at pre-amyloid stages in the 5xFAD mouse (Saminathan et al., 2026). This would suggest that Aβ plays an important role in the morphological and activity shifts identified in these microglia. Throughout the brain, regions with Aβ plaques contained altered microglia, identified visually as we could not analyze every region containing Aβ plaques as this model displays robust pathology in many regions.

Microglial morphology reflects their functional state, and changes in their morphology accompany disease, injury, infection, and aging (Wang et al., 2021). Microglia in visual brain regions burdened with Aβ plaques, the dLGN and V1, displayed amoeboid morphology as defined by a significant decrease in both microglial branch length and endpoints. Conversely, microglia in visual brain regions with insignificant Aβ plaque presence, the SC and SCN, displayed morphological characteristics similar to controls with longer branch length and more endpoints indicative of a ramified morphology. Amoeboid morphology, typically indicative of an inflammatory state, reveals that microglia in the aging dLGN and in V1 are activated. After identifying a morphological shift in microglia in specific brain regions, we aimed to determine the functional implications that reflect this shift. Microglia in the dLGN of 5xFAD mice show an increase in Iba1 and CD68 colocalization as they age, indicating they are increasingly phagocytic. We show that there is an increase in Aβ plaque coverage of dLGN which may explain the increasing phagocytic activity in the 5xFAD dLGN as these mice age. This may be due to an acute versus chronic response to Aβ plaque exposure. Reflective of the skeleton analysis, we see increased Iba1 and CD68 colocalization in regions with Aβ plaques, characteristic of phagocytic activity, but almost no phagocytic activity in regions devoid of Aβ plaques. These data point to phagocytic activity of amoeboid microglia in visual brain regions with significant Aβ plaque presence which is consistent with prior research showing that Aβ-associated microglia are phagocytic while tau-associated microglia lean toward a potentially neurotrophic function (Gerrits et al., 2021). The shift in microglial morphology occurs ahead of the phagocytic activity as we see in the 6-, 9-, and 12-month dLGN where morphologically, the microglia are amoeboid even at 6 mo while the phagocytic activity is not significantly increased until 9 mo. At 6 mo, there was not a significant increase in phagocytic activity in the 5xFAD mice compared to age-matched controls, but by 9 mo and 12 mo, we see a significant level of phagocytic activity compared to controls. This indicates that in the 5xFAD model, microglia are likely attempting to phagocytose Aβ plaques, and that the morphological shift leads to microglial activation.

Taken together, these results indicate that Aβ plaque deposition influences microglia morphology, and therefore activity, in visual brain regions of the 5xFAD mouse. In visual brain regions affected by AD-like pathology, specifically the presence of Aβ plaques, we see loss of microglial ramification and amoeboid microglial morphology as well as increased microglial labeling for the phagocytic marker CD68. Therefore, microglia in visual brain regions that present with significant levels of Aβ plaques, the dLGN and V1, are observed to have an amoeboid morphology and phagocytic function while regions that experience low or no Aβ plaque deposition, the SC and SCN, maintain their ramified microglial morphology and low to no phagocytic activity in the 5xFAD mouse model.

It is unclear as to why certain visual brain regions develop Aβ plaques but not others. It has been reported that parental origin of the transgene can alter Aβ plaque burden in these mice (Sasmita et al., 2025). This could also indicate there may be altered expression based on brain region (Gregg et al., 2010). We did not track the inheritance of this gene in our animals, and although this could be a confounding factor, it also allows for more biological variability within the colony better reflecting human genetics.

Disruption of the blood-brain barrier is a consequence of AD pathology partially induced by pathogenic Aβ isoforms (Petrushanko et al., 2023). As such, infiltrating macrophages have been shown to localize with Aβ plaques. In our study, we did not differentiate between infiltrating macrophages and microglia as both are labeled by Iba1 immunostaining. Although not a significant population, infiltrating macrophages may still be present in our analysis. Additional staining methods would be required such as using TMEM119, a microglia-specific marker, and CCR2, a marker of monocyte-derived macrophages in order to resolve these populations further (Fiala et al., 2002; Muñoz-Castro et al., 2023). Continued exploration of microglial activity would also reveal more specific behavior and function, such as using TREM2, P2Y12, or markers of lysosomal synapses to examine digestive exophagy as a potential mechanism of disease proliferation (Jacquet et al., 2024). The process of digestive exophagy, which involves microglia exocytosing amyloid fibrils toward Aβ plaques, may be a reason we see increased Aβ plaque size over time in the dLGN.

Future work should include an in-depth analysis of astrocytic morphology and function in these regions to determine whether they are contributing to clearance of Aβ plaques or leading to further neurodegeneration as they have been shown to cluster around Aβ plaques similar to microglia (Liu et al., 2024; Tsering et al., 2025). Continued work from our lab will compare these data to similar results collected from 3xTg-AD mice in order to determine effects of Aβ plaques as compared to tau tangles.

## Author Contributions

Conceptualization: SM, MJVH; Investigation: SM, JCS; Formal analysis: SM, JCS, MVH; Funding acquisition: SM, MJVH, Investigation: SM, JCS, MVH; Supervision: MVH; Writing – original draft: SM; Writing – review and editing: SM, JCS, MJVH.

## Data Availability Statement

The datasets generated for this study will be made available upon reasonable request.

